# Drug Resistance Mutations In Transplant Recipients With Suspected Resistance

**DOI:** 10.1101/2022.11.29.518463

**Authors:** Irene González, David Tarragó

## Abstract

Resistant CMV infections are challenging complications after SOT and HSCT. Prompt recognition of ARMs is imperative for appropriate therapy. 108 plasma samples from 96 CMV+ transplant recipients with suspected resistance were analysed in CNM in a retrospective nationwide study from January 2018 to July 2022 for resistance genotyping. ARMs in UL97 and UL54 were found in 26.87 % (18/67) and 10.60 % (7/66) of patients, respectively. Patients’ ARM distribution in UL97 was as follows: L595S n=3; L595S/M460I n=1; L595S/N510S n=1; L595W n=1; C603W n=4; A594V n=3; A594E n=1; C607Y n=1; L397R/T409M/H411L/M460I n=1; L397I n=1; H520Q n=1; four patients showed ARMs in UL54 as well (F412C n=1; T503I n=2; P522S n=1), whereas three patients exhibited ARMs in UL54 only (L501I/T503I/L516R/A834P n=1; A987G n=2). L516R in UL54 and L397R/I and H411L in UL97 have been found for the first time in a clinical sample. L595S/W was the most prevalent ARM found to lend resistance to GCV. In UL54 all ARMs lent resistance to GCV and CDV. In addition, A834P, found in one patient, also lent resistance to FOS. CMV load did not differ significantly in patients with or without ARMs, and no differences were found either between patients with ARMs in UL97 or in UL97 and UL54. Despite extensive use of classical antivirals for the treatment of CMV infection after HSCT and SOT, ARMs occurred mainly in viral UL97 kinase, which suggests that CDV and mostly FOS continue to be useful alternatives to nucleoside analogues after genotypic detection of ARMs.

## INTRODUCTION

CMV is a herpesvirus with a high worldwide prevalence; it causes serious complications in immunocompromised patients, particularly those who are recipients of hematopoietic progenitors (HSCT) or solid organ (SOT) (1, 2). The effects of CMV disease in these patients are responsible for high morbidity and mortality rates, as well as an increased risk of long-term graft loss (2–4).

The effectiveness of the preventive strategies currently used has managed to limit the incidence of the disease in the months following transplantation (4, 5). However, prolonged antiviral treatments increase the risk of selecting drug-resistance viral strains (2, 4, 6), which, added to the scarce therapeutic options, becomes challenging for the management of transplant recipients. Drug resistance, defined as a viral genetic alteration that decreases susceptibility to one or more antiviral drugs, should be suspected when CMV viremia fails to improve or continues to increase after two weeks of appropriately dosed and delivered antiviral therapy (7). Consequently, the need for genotypic analysis to detect resistance mutations during therapies is imperative. Prophylaxis with GCV IV or VGCV oral is the treatment of choice. FOS is often the first choice for the treatment of UL97-mutant ganciclovir-resistant CMV. A major concern with FOS is its high nephrotoxicity, as well as the alternative CDV. Approved in 2017 by the US Food and Drug Administration for the prevention of CMV in HSCT recipients (8, 9), a novel therapeutic alternative, such as letermovir, that do not have cross-resistance with current treatments has become a concern due to the rapid development of resistance mutations described recently (10). Mutations conferring resistance to LET are most commonly mapped to UL56. The rates of ARM in SOT patients is 5-12% depending on the group of patients studied but often is higher than 20% in patients with suspected ARM (11).

This study aimed to analyse the frequency of the appearance of mutations in UL97, UL54 and UL56 associated with antiviral resistance in clinical samples obtained from CMV+ transplant recipients with suspected resistant CMV to antivirals.

## RESULTS

### Analysis of Antiviral Resistance Mutations in UL97, UL54 and UL56

108 CMV positive PCR plasma from 96 transplanted patients yielded sequence data which enabled the analysis of at least one of the three genes of study. Studied genes UL54, UL97 and UL56 were fully characterized in 66, 67 and 96 CMV-positive patients, respectively. In 9 patients UL54 were characterized but not UL97. In other 10 patients UL97 were characterized but not UL54. In 20 patients, only UL56 could be fully analysed.

ARM was found in 21 transplant patients, 19 of them SOT recipients and 2 HSCT (Table 1). Regarding ARMs in UL97, 3 were cardiac transplant recipients, 2 liver transplant recipients, 6 lung transplant recipients, 6 kidney transplant recipients and 1 HSCT. Regarding ARM in UL54, 1 cardiac transplant recipient, 1 kidney transplant recipient, 4 lung transplant recipients and 1 HSCT. No ARM was found in UL56.

**Table 1:** ARM and CMV load in 21 SOT patients with suspicion of resistance to antivirals.

| Patient | Transplant | UL54 | ARM | UL56 | UL97 | ARM | CMV load (UI/mL) | Antiviral* |
| --- | --- | --- | --- | --- | --- | --- | --- | --- |
| 1 | SOT-C | R | F412C | S | R | C603W | 9,83X10<sup>3</sup> | GCV, FOS |
| 2 | SOT-C | S | - | S | R | L397R / T409M<br>/ H411L / M460I | 1,00X10<sup>5</sup> | GCV |
| 3 | SOT-K | S | - | S | R | A594V | 7,29X10<sup>4</sup> | GCV |
| 4 | SOT-L | S | - | S | R | C603W | 1,53X10<sup>3</sup> | GCV |
| 5 | SOT-K | S | - | S | R | L595S/N510S | 1,38X10<sup>4</sup> | GCV |
| 6 | SOT-K | S | - | S | R | L595S | 5,92X10<sup>4</sup> | GCV |
| 7 | HSCT | R | A987G | S | S | - | 8,74X10<sup>3</sup> | VGCV,<br>CDV |
| 8 | SOT-K | S | - | S | R | L595W | 4,12X10<sup>4</sup> | GCV |
| 9 | SOT-K | S | - | S | R | C607Y | 6,83X10<sup>3</sup> | GCV |
| 10 | SOT-H | S | - | S | R | H520Q | 3,75X10<sup>5</sup> | GCV |
| 11 | SOT-L | R | T503I | S | R | C603W | 3,75X10<sup>5</sup> | GCV, FOS |
| 12 | SOT-L | S | - | S | R | L397I | 2,65X10<sup>3</sup> | VGCV |
| 13 | SOT-L | S | - | S | R | L595S | 2,84X10<sup>3</sup> | GCV |
| 14 | SOT-L | R | P522S | S | R | M460I/L595S | 6,57X10<sup>3</sup> | GCV, FOS |
| 15 | SOT-H | S | - | S | R | A594V | 7,85X10<sup>3</sup> | VGCV, FOS |
| 16 | SOT-C | S | - | S | R | L595S | 7,50X10<sup>3</sup> | GCV |
| 17 | HSCT | S | - | S | R | A594E | 3,85X10<sup>3</sup> | VGCV |
| 18 | SOT-K | S | - | S | R | A594V | 1,45X10<sup>4</sup> | VGCV |
| 19 | SOT-L | R | **T503I** | S | R | **C603W** | 3,24X10<sup>3</sup> | VGCV, FOS |
| 20 | SOT-K | R | **L501I /**<br>**T503I /**<br>**L516R /**<br>**A834P** | S | S | - | 3,60X10<sup>3</sup> | GCV, FOS |
| 21 | SOT-L | R | **A987G** | S | S | - | 1,21X10<sup>4</sup> | GCV, CDV |

\*Antiviral therapy before ARM testing. In bold red ARMs previously described as selected under drug in vitro (23). In bold purple ARM not previously described (23). SOT-C =SOT heart, SOT-K= SOT Kidney, SOT-L= SOT Lung, SOT-H=SOT Hepatic, R= resistant, S= susceptible wild type

T503I was the most prevalent ARM in UL54 (3/7 patients), followed by A987G (2/7 patients) and L595S in UL97 (5/18 patients), followed by C603W (4/18 patients), A594V (3/18 patients), M460I (2/18 patients). L397I, L397R, T409M, H411L, H520Q, N510S, L595W and C607Y were found in one patient. Moreover, four patients developed ARMs simultaneously in UL54 (F412C 1; T503I 2; P522S 1), and in three patients ARM was detected in UL54 only (L501I; T503I; L516R; A834P). ARMs L397R and H411L in UL97 and L516R in UL54, which were previously described as obtained by drug selection in vitro, were found in two patients. L397I in UL97, which was detected in one cardiac recipient, has not been described before.

### Viral load and the presence of ARM

Viral loads for the 96 patients included in the study are shown in table 1 supplementary material and table 1 for the 21 patients with ARMs. No significant differences were found between the viral load of the samples with and without ARMs, either with ARMs only in UL97 and UL54-UL97 or UL54 only. A viral load threshold of 9.86 x10^3^ UI/mL was established to be able to analyse complete sequences with enough feasibility and accuracy to characterize ARMs in the three genes. Below this threshold, only UL56 was fully sequenced in all clinical samples.

### Polymorphism in UL54 DNA polymerase and UL97 kinase

The occurrence of polymorphism in UL54 is concentrated in specific positions, mostly in S655L and F669L, but other mutations were also found such as T885A, R792C and D898N and a duplication SS in the 585 position. Four patients exhibited D605E mutation in UL97, one of them together with ARM C603W. No polymorphism was found in UL56 sequences.

## DISCUSSION

In this study, we developed a genotypic method of amplification through PCR and Sanger sequencing to analyse ARM in the UL54, UL56 and UL97 genes in clinical samples from 96 transplant recipients with suspected resistance to antivirals. To date, this is the study with the highest number of patients conducted in Spain. Moreover, according to a recent review of Chou S. (23), we discovered a novel ARM “L397I” in UL97. Additionally, other three ARMs, such as L397R and H411L in UL97 and L516R in UL54, which were previously described as selected under drug in vitro, we detected directly in clinical samples (23). Interestingly, mutation at 397 position of UL97 confers resistance to maribavir despite this drug was not used in any patient.

Rate of resistance to antivirals was 26.87 % for UL97, whereas 5.97% developed combined resistance to UL97 and UL54, and 4.54% to UL54 only. This rate was close to the 27% detected in SOT patients through Sanger sequencing in a previous study conducted in Barcelona (11). Most ARMs were found in SOT patients, mainly in kidney and lung transplant recipients as described elsewhere (12, 13).

Most ARM was detected only in UL97 (14/21, 66.66%), indicating that the use of classical antivirals such as CDV and FOS, whose action mechanisms do not depend on UL97 kinase, is a reliable therapeutic option despite their wide use in transplant patients as alternative drugs. There was involvement of both UL97 and UL54 in 19.04 % (4/21) of patients with ARM. Surprisingly, in three patients ARM was only found in UL54; this fact may be explained by the fact that some ARM in UL97 may have reverted to wild-type after switching therapy to FOS or CDV. In this sense, previous experiments have shown that, fortunately, the most common ARM found, L595S/W, reverts after a while, provided that the selective pressure of GCV is removed (14), suggesting a certain disadvantage of this ARM compared to susceptible wild-type. Of note is the high proportion of patients with treatment failure unrelated to ARM: 72,72% (48/66) and 89,23% (58/65) regarding UL97 and UL54, respectively. Unknown factors probably related to the patient’s condition and/or virus virulence may be also responsible for most refractory CMV infections. Therefore, the absence of response to treatment is not decisive to establish a case of antiviral resistance, and confirmation with genotypic methods (11, 23) is required at any rate (21). Only two HSCT patients had not refractory CMV infections, which is in agreement with previous studies indicating that resistant CMV infections remain a rare complication in HSCT recipients, whereas refractory infections are more commonly found (15).

In this study, we searched for consensus ARM related to the lack of effectiveness of the main antivirals used against CMV (GCV, FOS, CDV, VGCV and LET) (Table 1). The presence of each of the mutations can affect a single drug or several ones simultaneously. Among the mutations found in the UL97 gene, H520Q/E and C603W/R/S were previously associated with high rates of resistance to GCV. However, the role of others, such as D605E, is controversial and, depending on the study, may be regarded as a resistance mutation or a variant of the natural sequence (16). Recent recombinant phenotypic experiments indicated that this mutation did not confer resistance to GCV (23). Therefore, we did not consider D605E, found in three patients, as an ARM.

Concerning resistance to LET, previously described ARMs were related to mutations located between amino acids 230 and 370 of UL56 (10, 15). In vitro and clinical studies showed that ARM developed faster than in UL97 and UL54, which is a reason for increasing concern among clinicians and virologists. Regarding UL56, since two naturally occurring sequence polymorphisms (L241P and R369S) were described to confer 160-fold and 38-fold reduced susceptibility to LET (17), respectively, we decided to study this gene despite only one patient with suspected resistance was treated with LET and, even with treatment failure, no ARM was found in UL56. Although the main target of ARM to LET has been found in UL56, other ARMs in UL51 and UL89 could not be ruled out. Seven patients with ARMs in UL54 were found, four of them with combined ARMs in UL97, which suggests that most of the ARMs were accumulated in UL97 kinase when GCV or a closely related antiviral as VGCV was used. This finding is in agreement with previous studies, in which more than 90% of ARMs occurred in the UL97 gene, specifically between codons 460-520 and 590-607 (3, 6, 13, 23). Other antivirals, such as FOS and CDV could be used instead in these cases, which highlights the importance of genotypic determination of ARMs for a right therapeutic choice. ARM was also found in UL54 DNA polymerase being T503I the most common (3/7 patients) which has been described as conferring resistance to GCV and CDV as well as A987G (2/7 patients). One patient developed multiple ARMs in UL54, one of which (A834P) is related to the appearance of resistance to FOS (18, 19).

In addition to the above-mentioned ARMs, other mutations compared to reference wild-type strains were found because of a certain polymorphism in UL54. The frequency of some of them is high, as in the case of S655L (51.14%) and F669L (42.86%) located at UL54. However, their consideration as candidate ARMs requires further recombinant phenotypic or marker transfer studies. It should be noted that the occurrence of multiple ARMs, which markedly increases antiviral resistance, thus complicating prognosis and treatment management (19, 20), was a common event: (8/21) of patients with ARMs.

In the search of ARMs in cohorts of patients with suspected resistance to antivirals, efforts have been made in many laboratories worldwide to develop NGS-based methods due to their ability to multiplex large numbers of samples. However, in our experience, for routine virological screening with few patients, NGS assays are still quite costly and time-consuming compared to PCR and Sanger sequencing. The main advantage of NGS was that ARMs may be characterised in samples with lower viral load (11) or when minor resistant subpopulations exist.

Despite limitations, the findings of this work contribute to reinforce the observation of the presence of mutations associated with drug resistance previously described, while making a case for the discussion on the involvement of new ones in the emergence of antiviral resistance. It is also shown that drug resistance is an important feature of CMV pathogenesis in transplant recipients that may threaten transplant outcomes, while the value of genotypic testing to identify potential antiviral resistance mutations is highlighted, which in turn could contribute to a better virological diagnosis and clinical performance.

## MATERIALS AND METHODS

### Clinical Samples and Transplant patients

In this retrospective study, 108 plasma samples from 96 transplant patients with suspicion of CMV resistant to antivirals were submitted to National Center for Microbiology (CNM) by hospitals all over the country from January 2018 to July 2022, to undergo genotypic analysis of antiviral resistance through sequencing of *ul54* and *ul97* genes. Residual samples were stored at −80°C until genotypic LET resistance characterization through *ul56* gene sequencing was performed. Median age of patients was 56 years-old. 64 SOT patients (39 SOT-K, 11 SOT-H, 7 SOT-C, 7 SOT-L) received prophylaxis and 32 HSCT patients received pre-emptive therapy. Individual therapy, viral load, gender, age and region where patient was living is detailed in Table 1 of supplementary material. Resistant and refractory CMV infection definitions were in agreement with consistent criteria (7). This study was approved by the Ethics Committee of the “Institute de Salud Carlos III” (CEI PI 11_2021-v3).

### DNA extraction, PCR design and sequencing

DNA extraction was performed from 200 μM of clinical sample (one sample per patient), using the “QIAamp Min ELUTE Virus Spin” Kit (QIAGEN), as per the manufacturer’s instructions. Systematic search and alignment of partial and complete sequences for the genes *ul54, ul56* and *ul97* were downloaded from GenBank database. Alignments using SeqMan (DNASTAR, Lasergen INC) and Mega X were performed to obtain the consensus and majority sequences, which were used as wild sensitive or resistance reference sequences. Three synthesized DNA fragments, containing all consensus resistance mutations described to date for each gene (22, 23) were cloned in E. coli plasmids and used as PCR and sequence-positive controls (Table 2). Three pairs of oligonucleotides were designed for PCR amplification of 990, 2246 and 649 bp fragments from ul97, ul54 and ul56, respectively. In addition, eight for UL54, six for UL56 and six for UL97 oligonucleotides were designed for Sanger sequencing (Table 3). Reactions were performed in Biorad C1000 Touch Thermal Cycler in a volume of 50μL and using Platinum SuperFi II DNA Polymerase (Thermo Fisher, Invitrogen), according to the manufacturer’s instructions. The oligonucleotides used to carry out the amplification were at a final concentration of 0.9 μM. PCR conditions for each gene are detailed in Table 4.

**Table 2.**
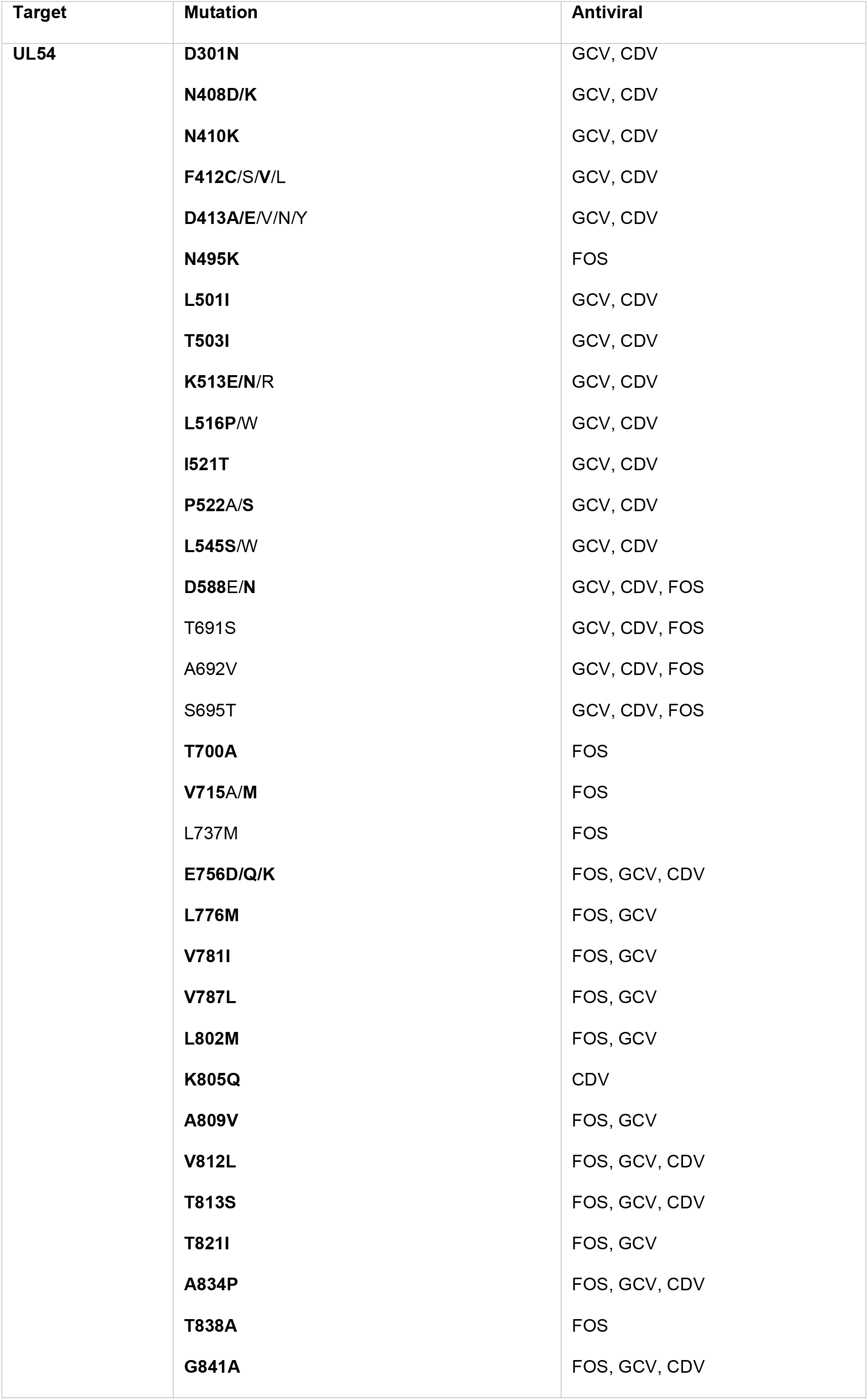
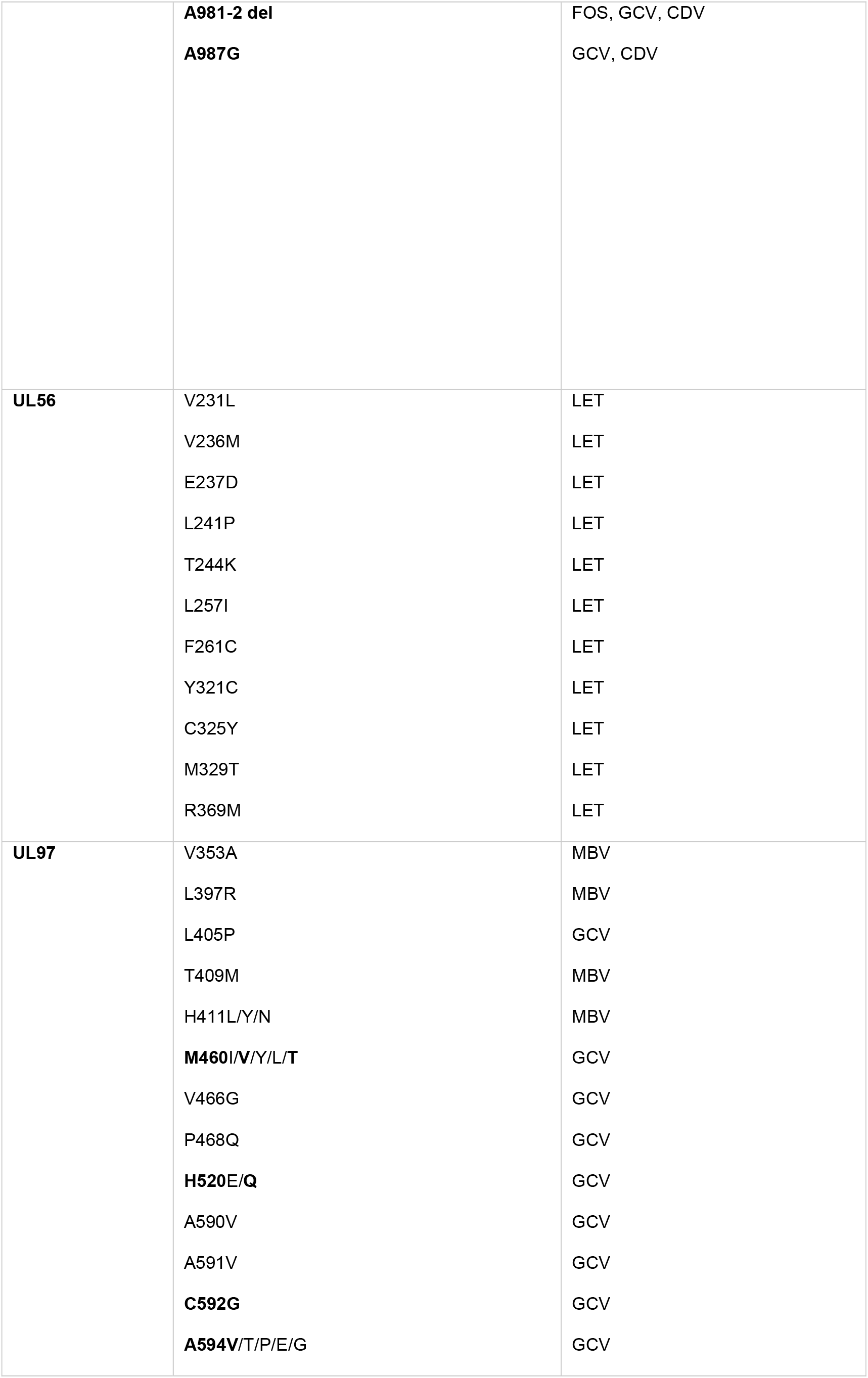
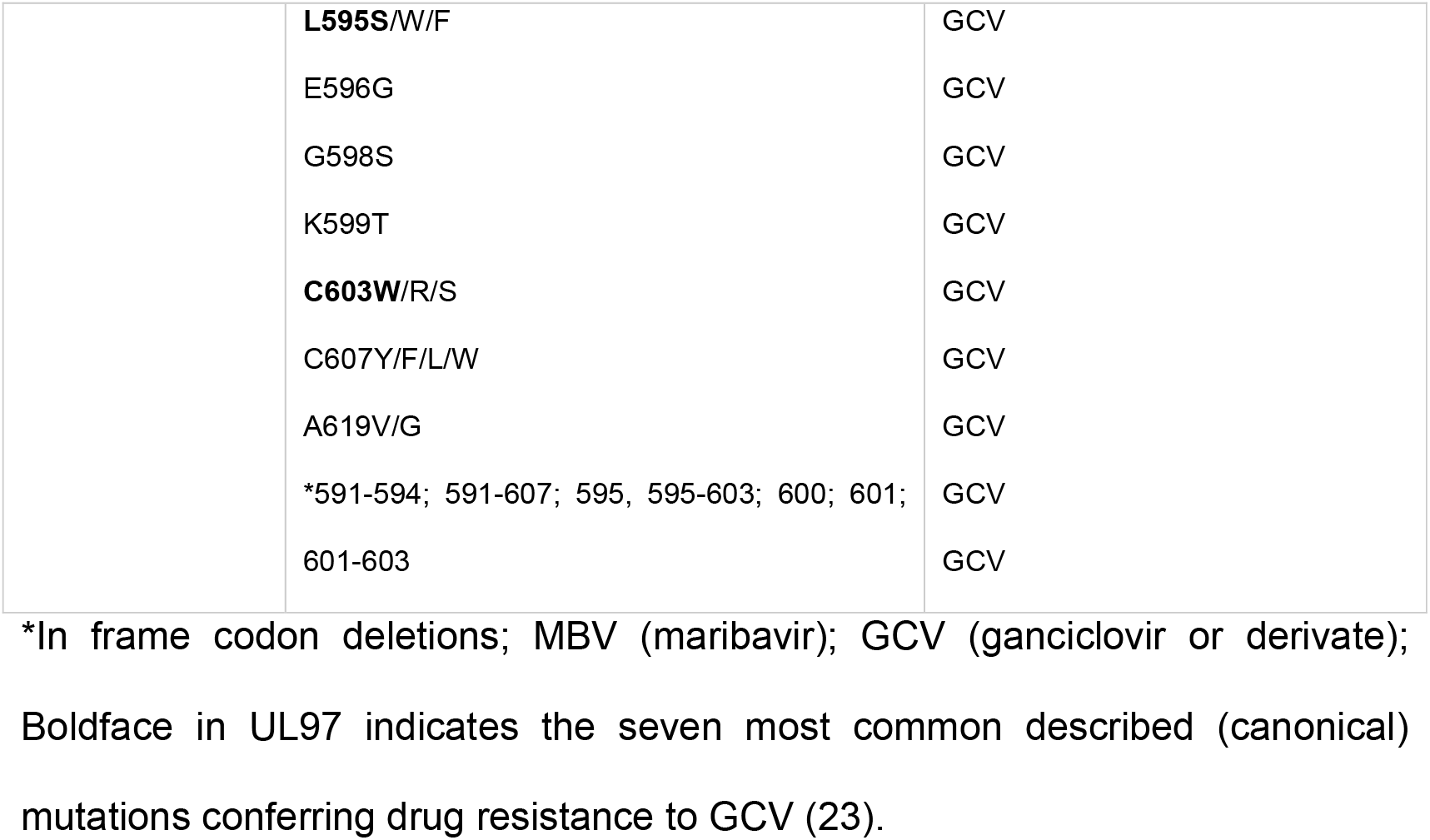
Previously described mutations associated to resistance to antivirals (23).

**Table 3.**
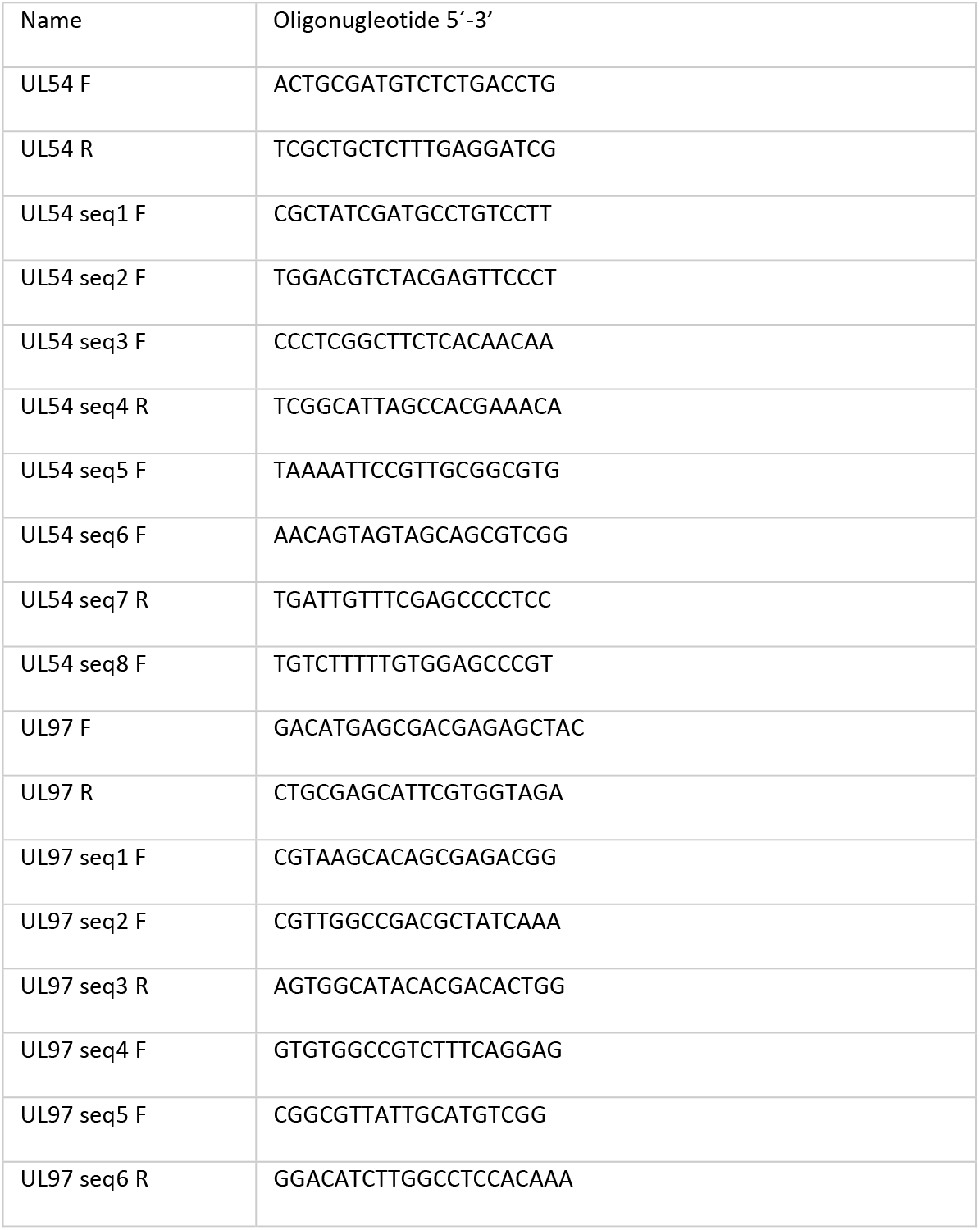
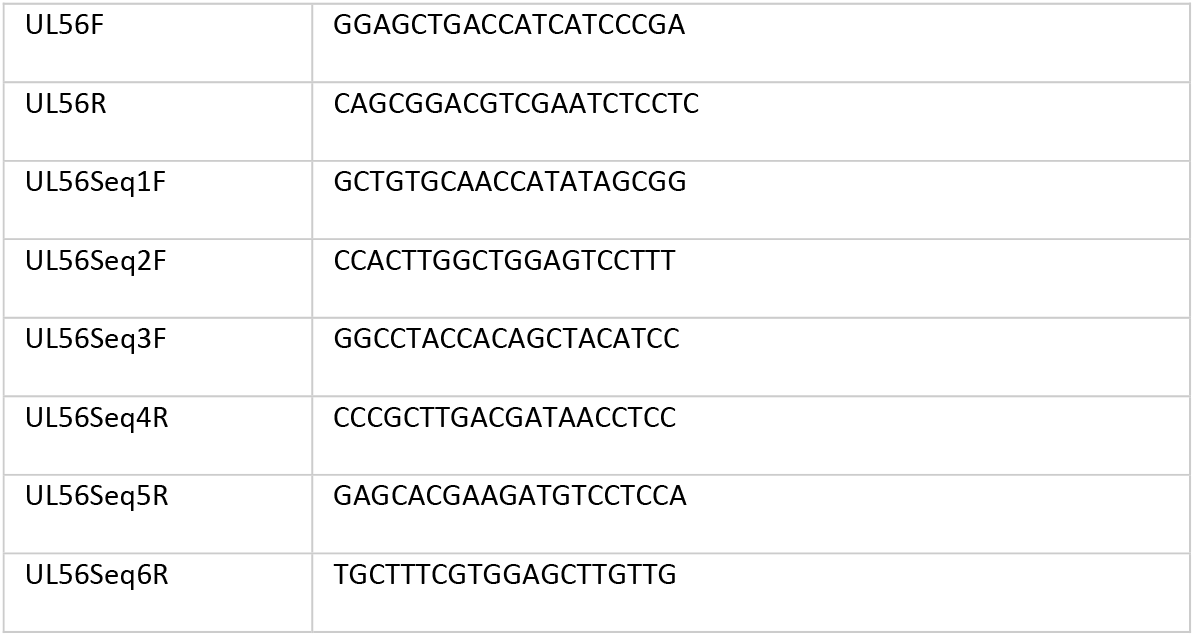
Oligonucleotides designed in the study for PCR and sequencing.

**Table 4.** PCR conditions for the amplification of target genes.

| Temperature |  |  |  | Time (min) |
| --- | --- | --- | --- | --- |
|  | <u>UL54</u> | <u>UL56</u> | <u>UL97</u> |  |
| Denaturation | 98 ° C | 98 ° C | 98 ° C | 00:30 |
| • Cycling 35x |  |  |  |  |
| Denaturation | 98 ° C | 98 ° C | 98 ° C | 00:10 |
| Annealling | 60 ° C | 60 ° C | 60 ° C | 00:15 |
| Extension | 68 ° C/1 min | 68 ° C/ 00:30 | 68 ° C/00:30 |  |
| Extension | 72 ° C | 72 ° C | 72 ° C | 05:00 |

A PCR product was considered available for sequencing when a detectable band of appropriate molecular weight was obtained by electrophoresis. Pre-sequencing purification of the PCR product was performed with the ExoProStarTM Enzymatic PCR and Sequence Reaction Clean-Up Kit 500 reactions (IllustraTM, Germany), following the manufacturer’s instructions. PCR products were processed for Sanger dideoxy sequencing with BigDye v. 3.1 (Applied Biosystems) in ABI PRISM 3100 sequencer (Applied Biosystems, California, USA).

### Multiplex real-time PCR for determination of CMV and EBV viral load and detection of HHV6, HHV7 and HHV8

We developed a 6-plex real-time PCR assay that is currently used in Reference Laboratory for Immune Preventable Diseases of National Centre for Microbiology. It was able to detect HHV6, HHV7 and HHV8 and to detect and quantify CMV and EBV. Quantitation used two sets of quantitative standards (for CMV and EBV) produced as follows: Relevant fragments of DNA (those amplified in real-time PCR) were inserted in a plasmid and cloned in transformed E. coli. Extracted serial dilutions of DNA from culture media were standardized against reference WHO material provided by Health Protection Agency (UK) for determination of CMV and EBV viral load. This multiplex assay included plasmid DNA positive control for HHV6, HHV7, HHV8 and an internal control (IC) of amplification. CMV/EBV quantitation demonstrated a sensitivity of 10 UI/mL and a wide dynamic range between 10-106 IU/mL for quantification of CMV and EBV in clinical samples and detection of HHV6, HHV7, HHV8 and an IC simultaneously. Quantitation accuracy was assessed with 2013 Cytomegalovirus and Epstein-Barr (DNA) EQA panels of QCMD and it was checked yearly using WHO standards. Primers (Sygma) and probes (Metabion) are in Table 5.

**Table 5:**
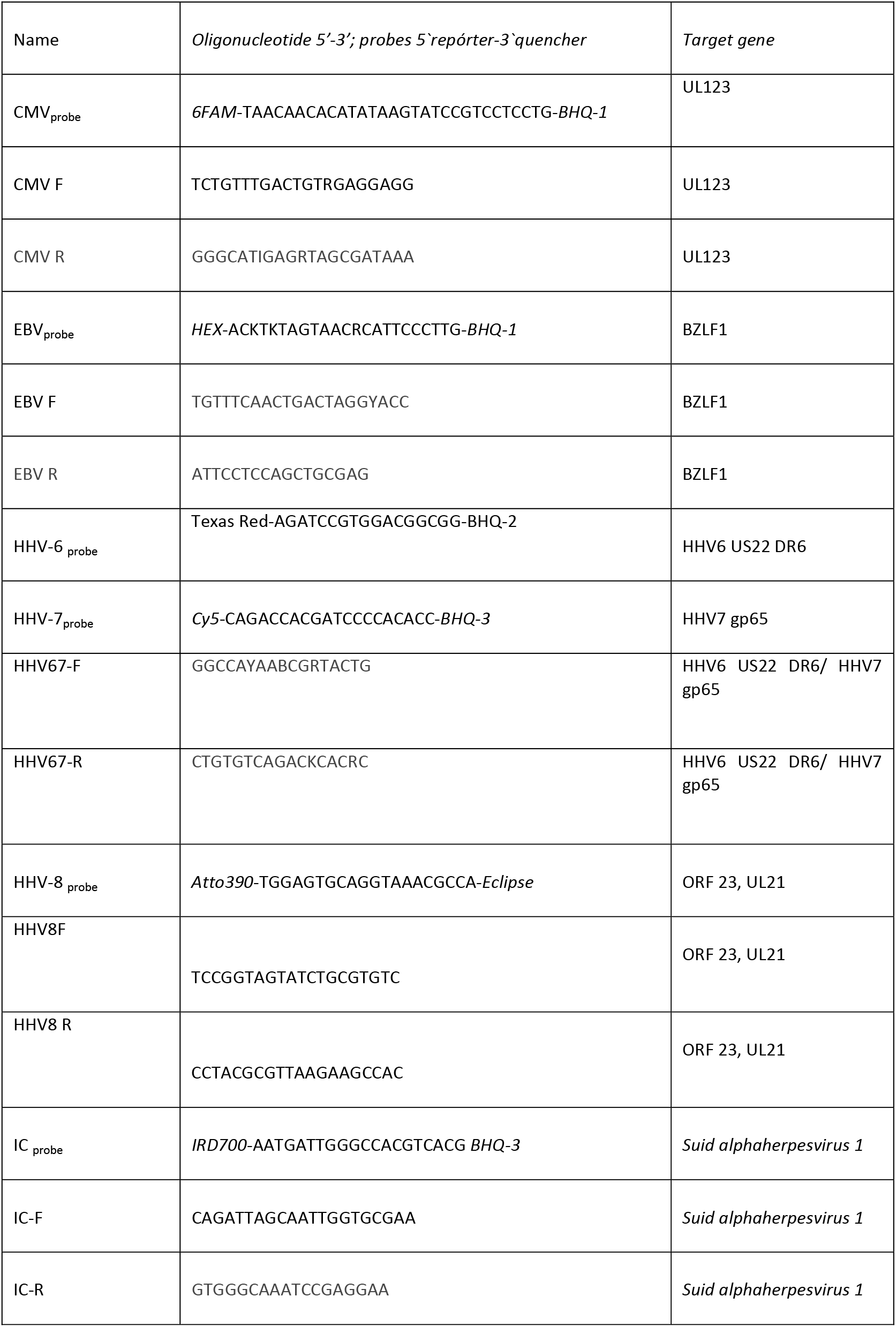
Primers and probes used in Multiplex real time PCR.

Amplification was carried out in a Rotor Gene thermocycler 6-plex with Quantitect Multiplex PCR kit (Qiagen) with 0,24μM of each primer y 0,25μM of each probe under the following conditions: Hold 95°C 15 min; 6 cycles of 94°C 30sec, 61°C 30sec; 40 cycles of 95°C 20sec,58° 60sec; hold 40°C 2 min.

### Sequence data and statistical analysis

The analysis and editing of the sequences was carried out with the SeqMan (Lasergene) and MegaX software. Amino acid sequences obtained were included in a database with previously created sequences containing all described ARMs for feasible searching of resistance mutations as well as sequences from reference laboratory strains such as Towne and AD169. Statistical analysis was performed using SPSS v28.0 software (SPSS, Chicago, IL). One-Way ANOVA was used to compare the viral load of CMV between clinical samples with and without ARMs and between clinical samples with ARMs in UL97 only and in UL54 plus UL97, followed by Tukey’s test for multiple comparisons between means (SD), 95 % CI and p-value ≤ 0.05.

### Limitations of the study

CNM service portfolio includes characterization of resistance mutations in UL97 and UL54. Treatment with LET was carried out in only one patient. However, due to the rapid emergence of ARMs in UL56, its characterization was included to know if a basal level of ARM occurred. Sanger sequencing is not able to detect subpopulations of CMV below 20-30% of the total, therefore minor subpopulations of CMV with ARMs, if any, were not identified. We established that direct amplification of clinical samples and sequencing required a viral load threshold ranging from 10^3^ UI/mL to 10^4^ UI/mL in order to obtain high-quality sequences for feasible analysis. In contrast, real-time PCR was able to detect below 10^2^ UI/mL. Some relevant characteristics of patients such as CMV serostatus (D/R) or days after SOT or HSCT were not available.

## Abbreviations

CMV: cytomegalovirus
EBV: Epstein-Barr Virus
HHV-6: human herpesvirus 6
HHV-7: human herpesvirus 7
HHV-8: human herpesvirus 8
CNM: National Center for Microbiology
CMV+: positive CMV real-time PCR
SOT: solid organ transplantation
HSCT: hematopoietic stem cells transplantation
ARM: antiviral resistance mutation
LET: Letermovir
FOS: Foscarnet
GCV: Ganciclovir
CDV: Cidofovir
VGCV: Valganciclovir

*Data availability including additional information regarding clinical samples and patients is in table 1 of Supplementary material.

## Acknowledgments

We are very grateful to Sergi Arteaga for kindly preparing manuscript.

This work was supported by a grant from Instituto de Salud Carlos III. AESI2021 PCIII00011-MPY434/2021. The funder had no role in study design, data collection and analysis, decision to publish, or preparation of the manuscript.

